# d-PBWT: dynamic positional Burrows-Wheeler transform

**DOI:** 10.1101/2020.01.14.906487

**Authors:** Ahsan Sanaullah, Degui Zhi, Shaojie Zhang

## Abstract

Durbin’s PBWT, a scalable data structure for haplotype matching, has been successfully applied to identical by descent (IBD) segment identification and genotype imputation. Once the PBWT of a haplotype panel is constructed, it supports efficient retrieval of all shared long segments among all individuals (long matches) and efficient query between an external haplotype and the panel. However, the standard PBWT is an array-based static data structure and does not support dynamic updates of the panel. Here, we generalize the static PBWT to a dynamic data structure, d-PBWT, where the reverse prefix sorting at each position is represented by linked lists. We developed efficient algorithms for insertion and deletion of individual haplotypes. In addition, we verified that d-PBWT can support all algorithms of PBWT. In doing so, we systematically investigated variations of set maximal match and long match query algorithms: while they all have average case time complexity independent of database size, they have different worst case complexities, linear time complexity with the size of the genome, and dependency on additional data structures.

## 1 Introduction

Durbin’s positional Burrows-Wheeler transform (PBWT) [2] is a scalable foundational data structure for modeling population haplotype sequences. It offers efficient algorithms for matching haplotypes that approach theoretically optimal complexity. Indeed, PBWT has been applied to important tasks such as genotype imputation [4], identification of identical by descent (IBD) segments [6], and genealogical search [5]. This has produced methods that scale to biobank scale datasets. The original PBWT paper described an array version of the PBWT, and a set of basic algorithms: Algorithms 1 and 2 for construction, Algorithms 3 and 4 for reporting all versus all long matches and set maximal matches, and Algorithm 5 for reporting set maximal matches between an out-of-panel query against a constructed PBWT panel. Recently, Naseri *et al.* [5] presented a new algorithm, L-PBWT-Query, that reports all long matches between an out-of-panel query against a constructed PBWT panel in time complexity linear to the length of the haplotypes and constant to the size of the panel. Naseri *et al.* introduced Linked Equal/Alternating Positions (LEAP) arrays, an additional data structure allowing direct jumping to boundaries of matching blocks. This algorithm offers efficient long matches, a more practical target for genealogical search. Arguably, L-PBWT-Query makes PBWT search more practical as it returns all long enough matches rather than merely the best matching ones. We believe that L-PBWT-Query represents a missing piece of the PBWT algorithms.

However, all above algorithms are based on arrays, which do not support dynamic updates. That means, if new haplotypes are to be added to, or some haplotypes are to be deleted from an existing PBWT data structure, one has to rebuild the entire PBWT, an expensive effort linear to the number of haplotypes. This will be inefficient for large databases hosting millions of haplotypes as they may face constant update requests per changing consent of data donors. Moreover, lack of dynamic updates prohibits PBWT to be applied to large-scale genotype imputation and phasing, which typically go through the panel multiple times and update individual’s haplotypes in turn. It is much more efficient to allow updating the PBWT with an individual’s new haplotypes while keeping others intact.

In this work we introduced d-PBWT, a dynamic version of the PBWT data structure. At each position *k*, instead of keeping track of sequence order using an array, we use a linked list, whose nodes encapsulate all pointers needed for traversing PBWT data structures. Our main results are: We developed efficient insertion and deletion algorithms that dynamically update all PBWT data structures (Algorithms 1 and 4). In addition, we established that d-PBWT can do Durbin’s Algorithms 1–5 and L-PBWT-Query with the same time complexity as the static version PBWT. While Durbin’s Algorithm 5 and L-PBWT-Query are practically independent of the number of haplotypes in the average case, we found that they are not in the worst case. We show two search algorithms for set maximal matches and long matches with worst case linear time complexity, but requiring multiple passes (Algorithms 2 and 3), and one search algorithm for long matches with average case linear time complexity with single pass without additional LEAP arrays data structures (Algorithm 7). These three new search algorithms can also be applied to the static PBWT. Table 1 summarizes the major contributions of this paper.

**Table 1.** Summary of Algorithms on PBWT and d-PBWT

|  | Data Structure | Functions | Algorithm ID | Time Complexity and Comments |
| --- | --- | --- | --- | --- |
| Durbin [2] | PBWT | Construction | Alg. 1&2 | $O(MN)$ |
| | | All vs. All Long Matches and Set Maximal Matches | Alg. 3&4 | $O(MN)$ |
| | | Set Maximal Match Query | Alg. 5 | Avg. $O(N)^*$ |
| Naseri *et al.* [5] | PBWT | Long Match Query | Alg. 2<br>(L-PBWT-Query) | Avg. $O(N)^*$. One sweep with LEAP arrays. |
| **This work** | d-PBWT | Insertion | Alg. 1 | Avg. $O(N)$ |
| | | Deletion | Alg. 4 | Avg. $O(N)$ |
| | | Set Maximal Match Query | Alg. 2 | $O(N)$ |
| | | Long Match Query | Alg. 3 | $O(N)$. Three sweeps without LEAP arrays. |
| | | Long Match Query | Alg. 7 | Avg. $O(N)^*$. One sweep without LEAP arrays. |
| | | Construction | - | $O(MN)$. Use Durbin's Alg. 1&2 |
| | | All vs. All Long Matches and Set Maximal Matches | - | $O(MN)$. Use Durbin's Alg. 3&4. |
| | | Conversion | Alg. 5&6 | $O(MN)$ |

$M$ is the number of sequences. $N$ is the number of sites. $c$ is the number of matches found. Time complexities assume $c < N$ . The number of sweeps and LEAP arrays usage are only mentioned for long match query algorithms because they are important for genealogical search.
\* See Appendix A for discussion of the time complexity.

## 2 Methods

### 2.1 PBWT

The following is a review of Durbin’s PBWT paper and notation [2]. PBWT is a data structure that groups similar strings by sorting the reverse prefixes at each length. Say we have a PBWT data structure of a set *X* of *M* haplotype sequences *x*_*i*_ ∈ *X, i* ∈ {0 … *M* − 1}. Each sequence has *N* sites indexed by *k* ∈ {0 … *N* − 1}, values at a site are 0 or 1, *x*_*i*_[*k*] ∈ {0, 1}. For some haplotype sequence *s* we use *s*[*k*_1_, *k*_2_) to represent the substring of *s* beginning at *k*_1_ and ending at *k*_2_ − 1. The length of this substring is *k*_2_ − *k*_1_. Sequences *s* and *t* have a match from *k*_1_ to *k*_2_ if *s*[*k*_1_, *k*_2_) = *t*[*k*_1_*, k*_2_). This match is **locally maximal** if it can’t be extended, i.e., (*s*[*k*_1_ − 1] ≠ *t*[*k*_1_ − 1] **or***k*_1_ = 0) **and**(*s*[*k*_2_] ≠ *t*[*k*_2_] **or***k*_2_ = *N*). A match is a **long match** if it is locally maximal and at least length *L*. A match is a **set maximal** match from a sequence *s* to *X* if it is locally maximal and there is no longer match between *s* and any other sequence from *X* that covers the matching region.

The **prefix array** *a* contains *N*+1 sorted orderings of the sequences, one for each *k* ∈ {0 … *N*}. The *k*-th sorted ordering is *a*_*k*_, the ordering of *a*_*k*_ is based on the reversed prefixes *x*[0, *k*), if the prefixes are the same they are ordered according their index *i* in *X*. *a*_*k*_ can also be thought of as the sorted ordering of the reversed prefixes of length *k*. In any *a*_*k*_, adjacent sequences are maximally matching until *k*. *y*_*k*_ is the *i*-th sequence in 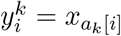. The **divergence array** keeps track of the start position of locally maximal matches ending at *k* between a sequence and the sequence above it in *a*_*k*_, i.e., *d*_*k*_[*i*] is the smallest value *j* such that 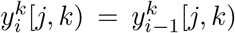. The **extension function***w*_*k*_(*i, h*), *h* ∈ {0, 1} gives the *a*_*k*+1_ index of the first sequence after *a*_*k*_[*i*] (*a*_*k*_[*i*] inclusive) that has *h* at site *k*, i.e.,

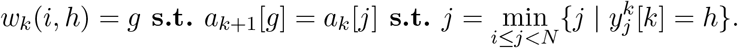

In implementation, the extension function is fully specified by two arrays: *u* and *v*. *w*_*k*_(*i,* 0) is stored at *u*_*k*_[*i*] and *w*_*k*_(*i,* 1) is stored at *v*_*k*_[*i*].

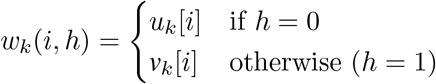

*v* is redefined here to make the extension function more intuitive.

### 2.2 d-PBWT

Our main observation is that PBWT algorithms are not necessarily array algorithms. The essence of PBWT is, at each site, sequences are ordered by their reverse prefix, and the updates of the ordering across adjacent sites tracked by pointers. However, the ordering of sequences is not necessarily tracked by prefix arrays. This fact might be not obvious as the original BWT [1] was based on arrays and all Durbin’s PBWT algorithms and previous PBWT algorithms are written in the array language. In this work we propose using a doubly linked list at each site to track the sorting. In doing so, we can enable PBWT for dynamic updates, while still maintaining all basic operations of PBWT. Below we formally describe the dynamic version of PBWT, d-PBWT, and all its algorithms.

Like PBWT, the d-PBWT consists of *N* columns^3^, each corresponds to one site^4^. Column *k* is a doubly linked list of *M* nodes that represents the reverse prefix sorting of all *M* sequences at site *k*. Each column has a top node. The top node of column *k* is noted as (*k,* 0), containing the first sequence in reverse prefix sorting at column *k*. A node *n* in column *k* is noted as (*k, i*) *iff* it takes *i* node traversals to reach *n* from the top node of column *k*. It turns out that we can encapsulate all necessary PBWT pointers at (*k, i*), including *a*_*k*_[*i*], *d*_*k*_[*i*], *u*_*k*_[*i*], and *v*_*k*_[*i*] inside individual nodes: A node *n* has one function, *w*, and six properties. The properties are *above, below,* sequenceID, *d, u,* and *v*. *n.above* represents (*k, i*−1) and *n.below* represents (*k, i*+1). *n*.sequenceID is an integer ∈ {0 … *M* − 1} that is unique to the sequence *n* represents, i.e., *n.*sequenceID is *a*_*k*_[*i*]. *n*[*j, k*) is equivalent to 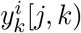 and *x*_*n.sequenceID*_[*j, k*). *n.d* is equivalent to *d*_*k*_[*i*], i.e., *n.d* = min_0≤*j≤k*_ **s.t.***n*[*j, k*) = *n.above*[*j, k*). Each node also has *u* and *v* pointers that make up the extension function, these are equivalent to the *u* and *v* arrays as well. This means that they point to the node in the next column of the first sequence below them (self included) that has 0 (for *u*) or 1 (for *v*). *n.w*(*h*) gets/sets *n.u* if *h* = 0, otherwise *n.v*. Lastly, the haplotype panel of d-PBWT is a dynamic array of *M* haplotypes. The equivalencies between data structures of PBWT and d-PBWT are summarized in Table 2.

**Table 2.** PBWT and d-PBWT equivalencies

| Entity | PBWT | d-PBWT |
| --- | --- | --- |
| Column $k$ | $a_k, d_k, u_k, v_k$ | linked list hung from top node at $k$ |
| Index $i$ at Column $k$ | $a_k[i], d_k[i], u_k[i], v_k[i]$ | node ($n$) that has $i$ nodes above it and is in column $k$ |
| Sequence name | $a_k[i]$ | $n.\text{sequenceID}$ |
| Match length | $d_k[i]$ | $n.d$ |
| Move to previous sequence in reverse prefix sorting | $a_k[i-1], d_k[i-1], u_k[i-1], v_k[i-1]$ | $n.\text{above}$ |
| Move to next sequence in reverse prefix sorting | $a_k[i+1], d_k[i+1], u_k[i+1], v_k[i+1]$ | $n.\text{below}$ |
| Move to next site by extension function | $u_k[i], v_k[i], w_k(i, h)$ | $n.u, n.v, n.w(h)$ |
| Substring in original haplotype | $y_k[j, k), x_{a_k[i]}[j, k)$ | $n[j, k), x_{n.\text{sequenceID}}[j, k)$ |

### 2.3 Insertion

The insertion algorithm works by first inserting the nodes of *z* in the correct position in each column and then calculating the divergence values after. This is analogous to first updating the prefix arrays and then updating the divergence arrays. This is done by first sweeping forwards through the data to insert the nodes, and then sweeping backwards through the data to calculate the divergence values. *z* is inserted into the dynamic haplotype panel in the forward sweep.

We update the prefix panel by keeping track of the node that *z* should be above and then inserting *z* above said node. We define *t*_*k*_ as the node that *z* should be above in column *k*. If we have *t*_*k*_, we can get *t*_*k*+1_ using the extension function. The sequence that will be below *z* at column *k* + 1 is the first sequence below *z* (not inclusive) that has the same value as *z* at *k*, i.e., *t*_*k*+1_ = *t*_*k*_.*w*(*z*[*k*]). We can use this to calculate all *t*_*k*_’s and insert *z* above them. See Figure 1.

**Fig. 1.**
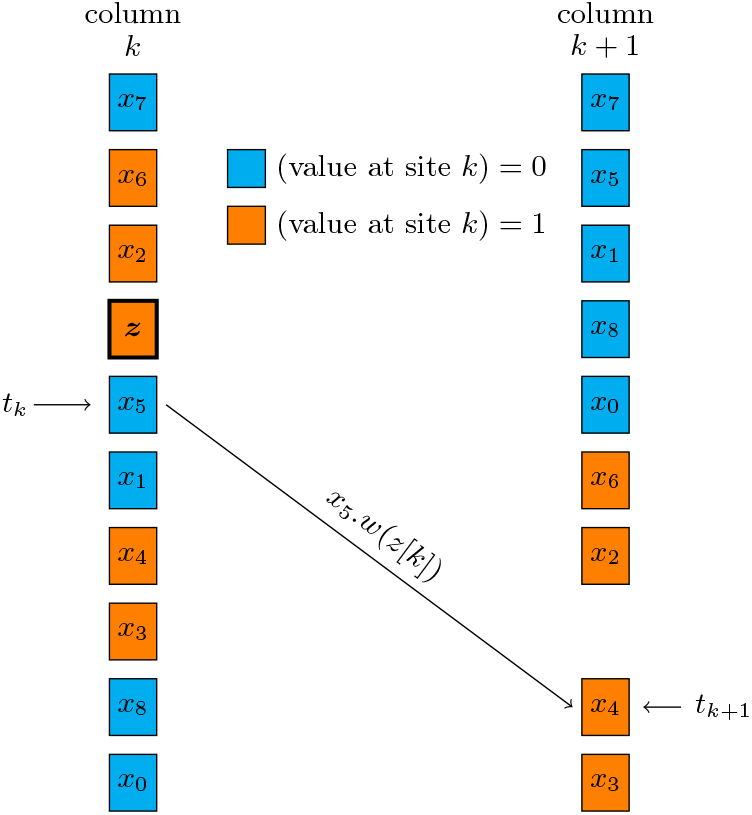
Finding the insertion location of *z* at next column. *t*_*k*_.*w*(*z*[*k*]) returns *t*_*k*+1_. This is because the first sequence below *x*_5_ (*t*_*k*_) that has the same value as *z* at *k* is *x*_4_. *t*_*k*_.*w*(*z*[*k*]) points to the *k* + 1 node of the first sequence below *t*_*k*_ (*t*_*k*_ inclusive) that has the same value as *z* at *k*. Therefore, *t*_*k*_.*w*(*z*[*k*]) points to the *k* + 1 node of *x*_4_.

We also have to maintain the *u* and *v* pointers. The contiguous group of sequences directly above *z* at *k* that have the opposite of *z*[*k*] at *k* need to have their 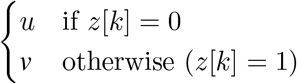 pointer updated to point to *z*_*k*+1_. Fortunately, because of *linkage disequilibrium* ^5^ this will be a small constant on average. Furthermore, *z*_*k*_.*w*(*z*[*k*]) is equal to *z*_*k*+1_ and *z*_*k*_.*w*(opposite of *z*[*k*]) is equivalent to *t*_*k*_.*w*(opposite of *z*[*k*]) (*z*_*k*_ is the node of *z* in colum *k*). Therefore *u* and *v* pointers of column *k* are updated after insertion of *z*_*k*+1_ into column *k* + 1. See Figure 2.

**Fig. 2.**
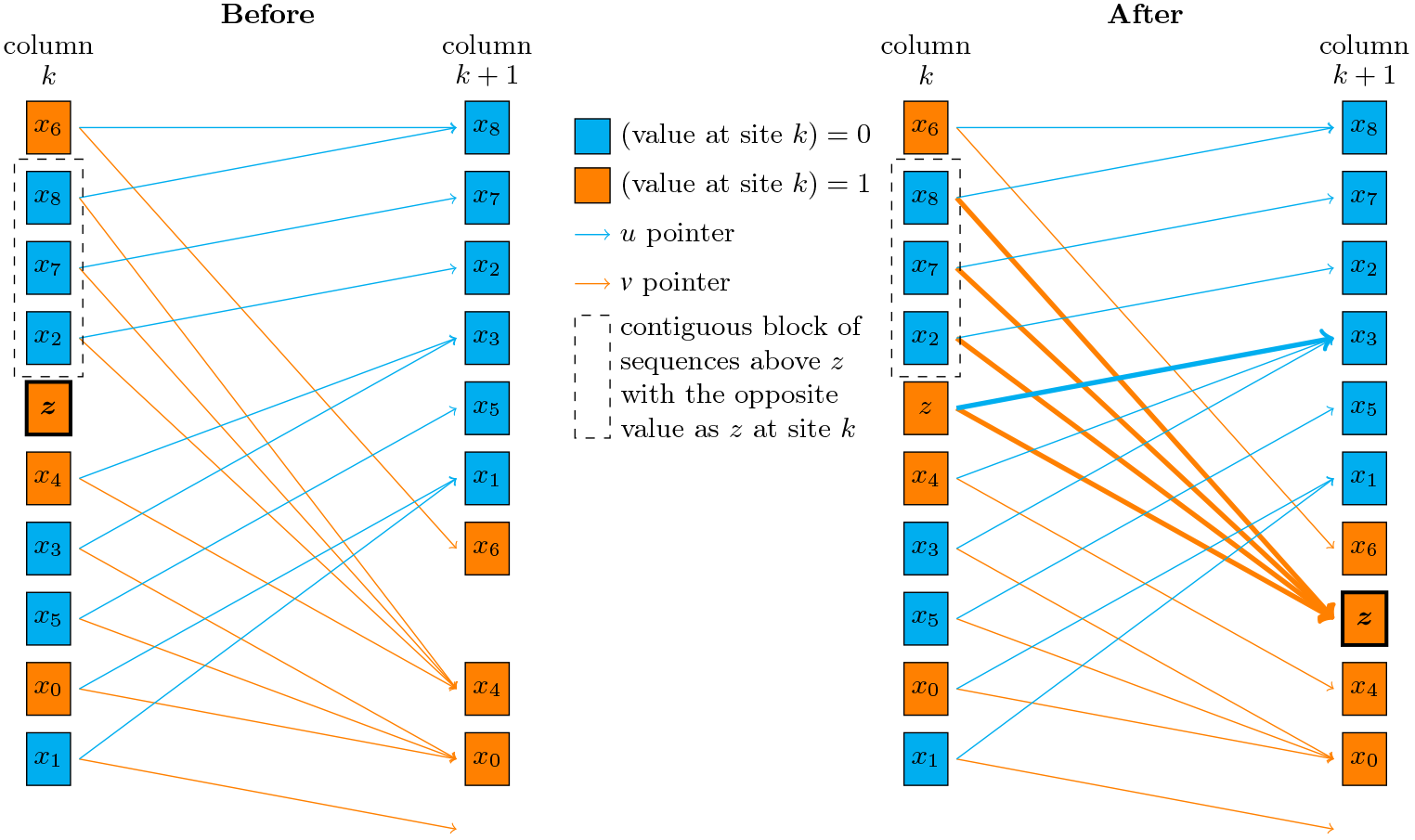
Updating the *u* and *v* pointers when inserting the *z* node into column *k* + 1. Updated items are bold, including *z*, four *v* pointers, and one *u* pointer. It also shows the update of the *w.*(opposite of *z*[*k*]) pointers of the contiguous block of sequences above *z* with the opposite value as *z* at site *k*.

The only thing left to do is update the divergence values. For each column *k*, only 2 divergence need to be set, the divergence of *z* and the divergence of *z.below*, all other divergence values remain unchanged because the sequence above all other sequences remain unchanged. We will update the divergence values by going backwards through the columns and keeping track of the minimum divergence value (longest match) found so far. A key observation is that at any column *k*, the divergence value of *z* must be at least the divergence value of *z* at *k* + 1, i.e., *z*_*k*_.*d* ≤ *z*_*k*+1_*.d*. This is true because if the sequence above *z* at *k* + 1 matches with *z r* sites backwards from *k* + 1, then that sequence will be above *z* at *k* and the sites will still match. The same goes for *z* and *z.below*.(See **Lemma 1** in Appendix B for the proof of this claim.) Therefore, to calculate the divergence values we go from *k* = *N* → 1 keeping track of divergence value of previous column. The divergence value of the *z*_*k*_.*below* and *z*_*k*_ is calculated by decrementing from divergence of *z*_*k*+1_*.below* and *z*_*k*+1_ until the first site that is different is found. See Figure 3.

**Fig. 3.**
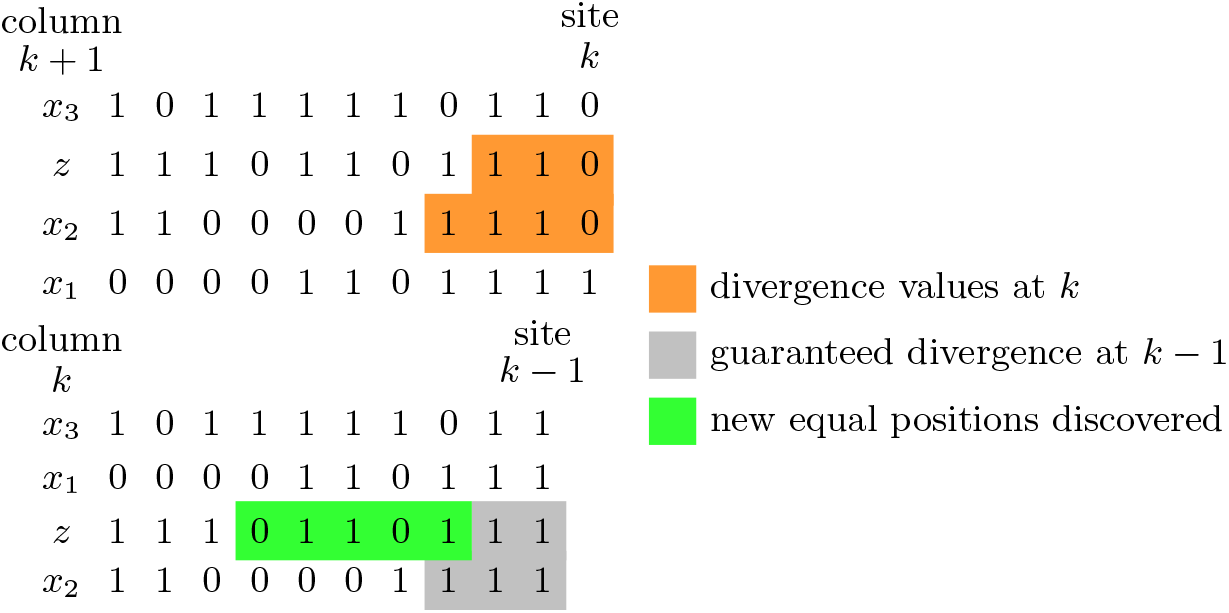
Updating the divergence value of *z* and *z.below* at position *k* based on position *k* + 1. At column *k* we know that there is some sequence above *z* that matches until the divergence value of *z* in column *k* + 1. This is because if the sequence is above *z* in column *k* + 1 and it matches at site *k*, then it is above *z* in column *k*. The relative order of sequences that have the same value at site *k* is the same in columns *k* and *k* + 1. The same goes for *z.below* and the divergence value of *z.below*.

The time complexity of the Insertion algorithm (Algorithm 1) is average case *O*(*N*). This is average case instead of worst case solely because of updating the *u* and *v* pointers and insertion of *z* into the dynamic haplotype panel. However, as stated, because of *linkage disequilibrium*, a case where the constant is non-negligible is extremely rare. Insertion of *z* into the dynamic haplotype panel is amortized *O*(*N*), therefore it is average case *O*(*N*). The insertion of the nodes of *z* into the correct position in each column is worst case *O*(*N*) because insertion into one column is constant time and there are *N* columns inserted into. The divergence calculation is also worst case *O*(*N*), because the outer loop runs for *N* iterations and the sum of all iterations of the inner loop will beat most *N*. The sum of all iterations of the inner loop will be at most *N* because it decrements an index from *N* → 0 over the whole algorithm. The fact that a “virtual insertion” algorithm (i.e., find all divergence and *t*_*k*_ values without updating *u* and *v* pointers or inserting *z*) is worst case *O*(*N*) will be used later to show the time complexity of the query algorithms.

**Algorithm 1:**
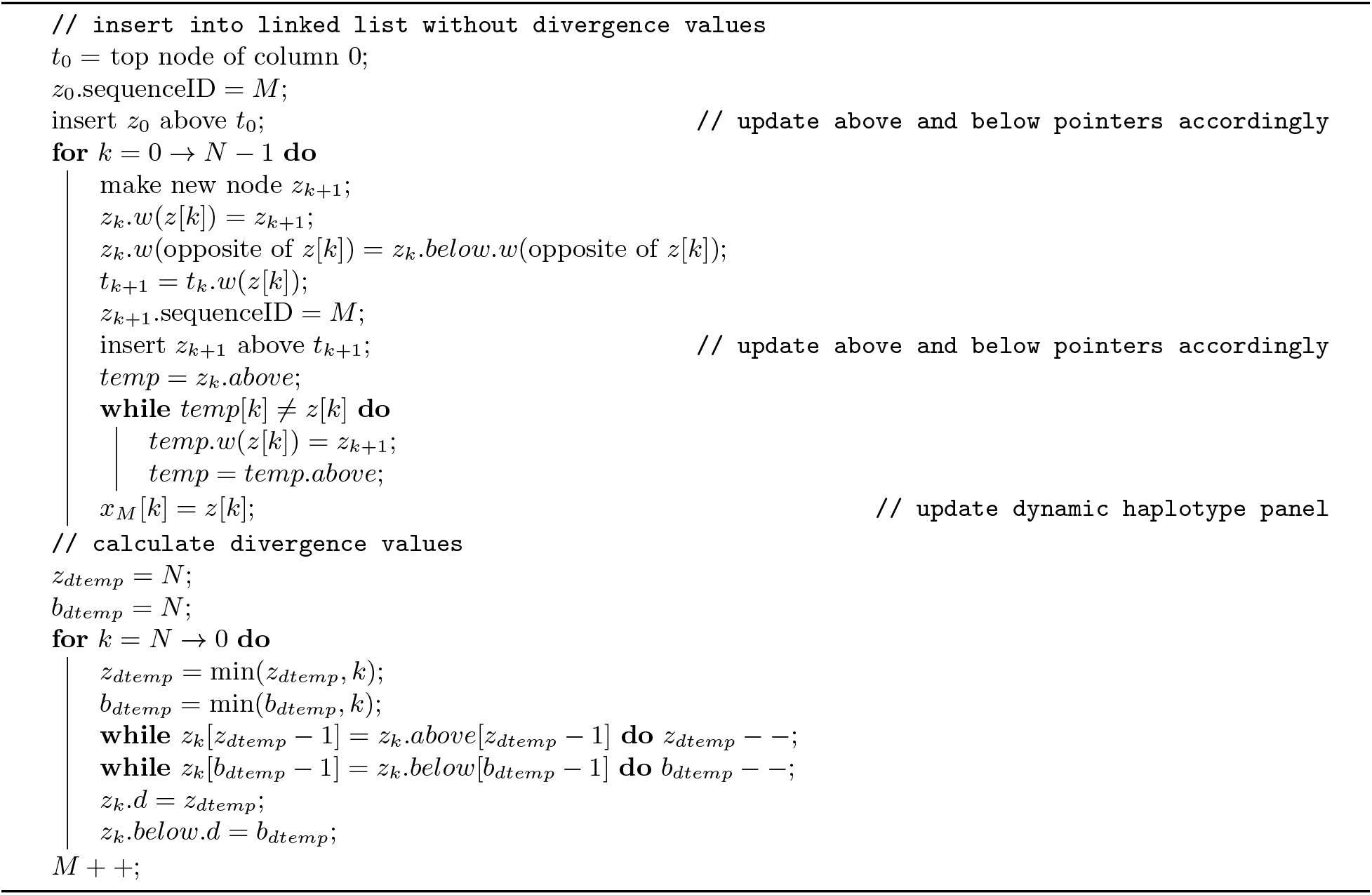
Insertion: Insert new sequence *z* into d-PBWT

### 2.4 Set Maximal Match Query

Durbin’s Algorithm 5 is not worst case *O*(*N*), refer to Appendix A for clarification. Nevertheless, we have empirical evidence of Algorithm 5’s *O*(*N*) performance in the average case [5]. Here we show a worst-case *O*(*N*) algorithm for outputting set maximal matches from *z* to *X*.

The set maximal match query virtually inserts *z* into the d-PBWT. The sweep back of the insertion algorithm is modified so that set-maximal matches are simultaneously outputted. The set maximal match query is fairly straightforward after one vital element is understood. If *z*’s locally maximal match ending at *k* matches farther back than its locally maximal match ending at *k* + 1, then *z*’s locally maximal match ending at *k* is a set maximal match (see **Lemma 2** in Appendix B for the proof). Therefore, we can just compare divergence values at *k* and *k* + 1 when calculating them to find and output set maximal matches.

A match is a set maximal if the match is locally maximal and there is no match with *z* that encompasses this match. We know that the match is locally maximal because we defined it as “locally maximal match ending at *k*” **and** it ends at *k*, therefore it is locally maximal (we know it ends at *k* because if it didn’t the locally maximal matches of *k* and *k* + 1 would match to the same point). Lastly, if there was a match with *z* that encompassed this match, then the locally maximal matches of *k* and *k* + 1 would match to the same point. Therefore, the *z*’s locally maximal match ending at *k* is a set-maximal match and can be outputted. Furthermore, there might be multiple sequences with this match, this is easily checked with divergence values.

**Algorithm 2:**
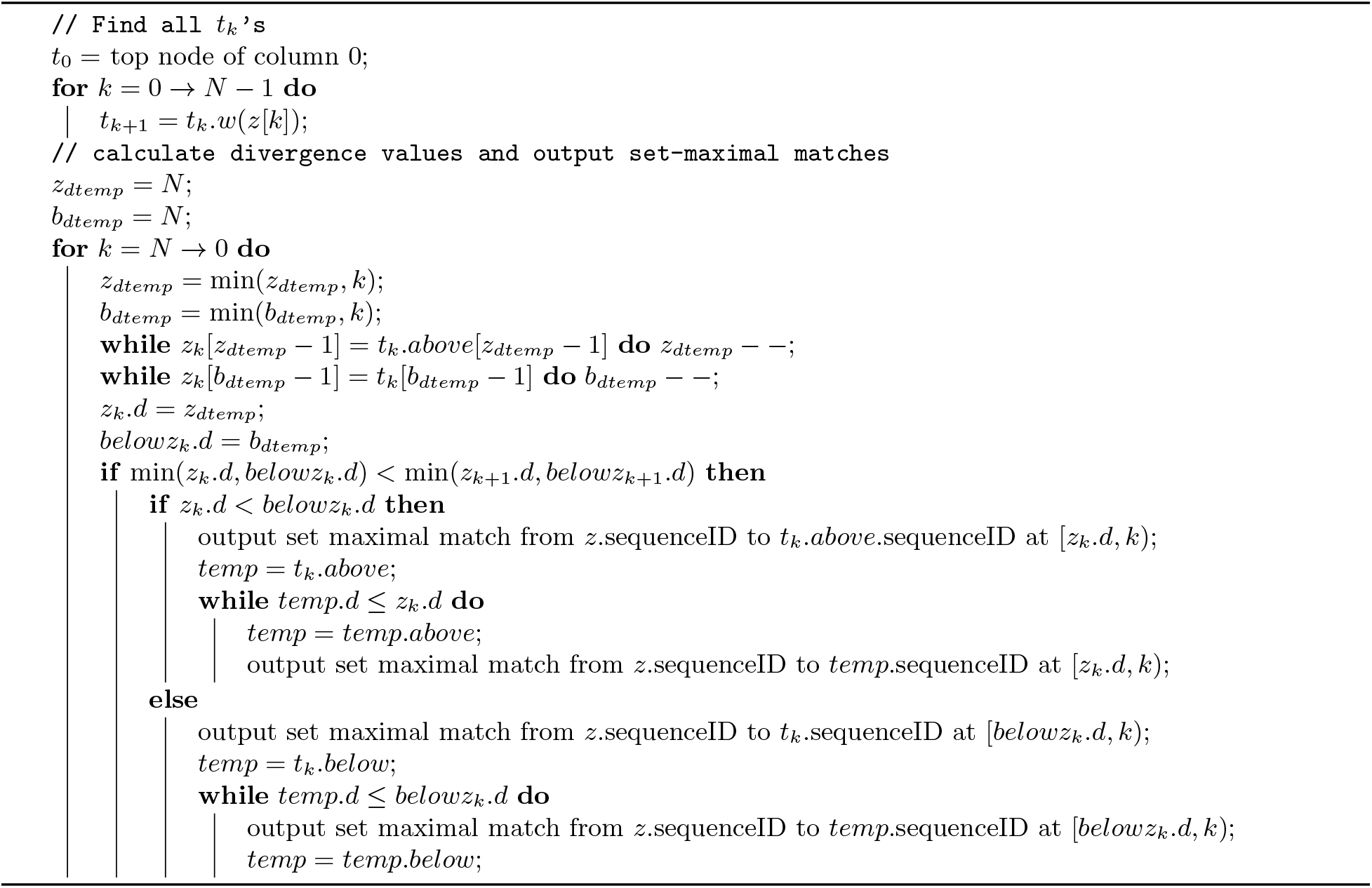
Set Maximal Match Query: Find set maximal matches from *z* to sequences in the d-PBWT

Lastly, since the sequence above and below *z* can’t match *z* with the same divergence, locally maximal matches will either be all above or all below, therefore only the direction with the smaller divergence value (longer match) will be checked. Assume the sequence above and below *z* in the sort order match *z* with the same divergence, the sequence above has value 0 one position behind and the sequence below has value 1. *z* must have either 0 or 1 at this position. Therefore the sequences above and below *z* do not match *z* with the same divergence.

The time complexity of the Set Maximal Match Query algorithm (Algorithm 2) is worst case *O*(*N* + *c*). The virtual insertion is *O*(*N*) because the haplotype panel and the *u* and *v* pointers are not updated. The while loops are only entered when there is a set maximal match to output and each match is outputted exactly once. Therefore the sum of iterations of the output while loops is bounded by *c* (number of matches found) and the whole algorithm is *O*(*N* + *c*).

### 2.5 Long Match Query

Naseri *et al.* [5] first proposed an efficient algorithm (L-PBWT-Query) to find all long matches between a query haplotype *z* and a database of haplotypes *X* in average case *O*(*N* + *c*) time by using PBWT and LEAP arrays to skip unnecessary checks (L-PBWT-Query is average case because it relies on Durbin’s Algorithm 5, see Appendix A). Here we propose a new algorithm for finding long matches without using LEAP arrays in d-PBWT in worst case *O*(*N* + *c*) time.

We will need the divergence values for our query algorithm, therefore the first thing we do is virtually insert *z* into the data structure. This means we get all the *t*_*k*_ values and all the new divergence values if *z* was inserted. Then we do a third sweep of the data while keeping track of a matching block. Note that we don’t update the haplotype panel or *u* and *v* pointers.

The high level idea of this algorithm is to keep track of the block of sequences that match with *z* length *L* or longer until *k*. We will denote the boundaries of this block 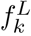 and 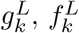 points to the first sequence in the block and 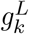 points to the first sequence below 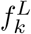 not in the block. Note that the definition of *f*^*L*^ and *g*^*L*^ is different from Durbin’s definition of *f* and *g*.

Given 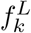 and 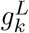, we want to get 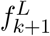 and 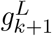. First, we use the extension function. 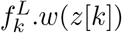 will give us the position in column *k* + 1 of the first sequence after 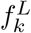 that has the same value as *z* at *k*. Likewise with 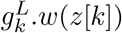. Therefore, 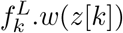 and 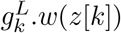 will give us the 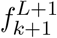 and 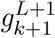 (see **Lemma 3** in Appendix B for the proof). 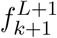 and 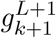 mark the boundaries of the block of sequences that match with *z* length *L* + 1 or longer until *k* + 1. The difference between 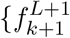, 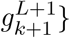 and 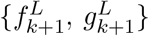 is only the sequences that match with *z* length *L* until *k* + 1. Therefore, we can intuitively use the divergence values to check if a sequence on the boundary matches with *z* length *L*, if it does, we move the boundary to include it in the block. After both boundaries reach a sequence that doesn’t match with *z* length *L* until *k* + 1, we have found 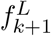 and 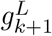.

There are two cases when we try to expand our 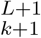 block. If it is not empty 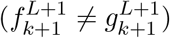, we can use the divergence values of the sequences in the d-PBWT to expand the boundaries. However, if it is empty 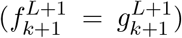, we must use the divergence values we calculated during virtual insertion to expand the boundaries initially. Lastly, when we expand the boundaries to include a new sequence in the block we also remember the starting position of the match (*k* + 1 − *L*) in an array *dZ* so that we can output it later. Meanwhile, 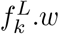(opposite of *z*[*k*]) and 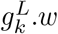(opposite of *z*[*k*]) will give us the block of sequences (in column *k* + 1) that have matches length *L* or longer until *k* and their match ends at *k*, we output these. We can repeat this procedure for all *k* to output all matches longer than *L* between query sequence and database. See Figure 4.

**Fig. 4.**
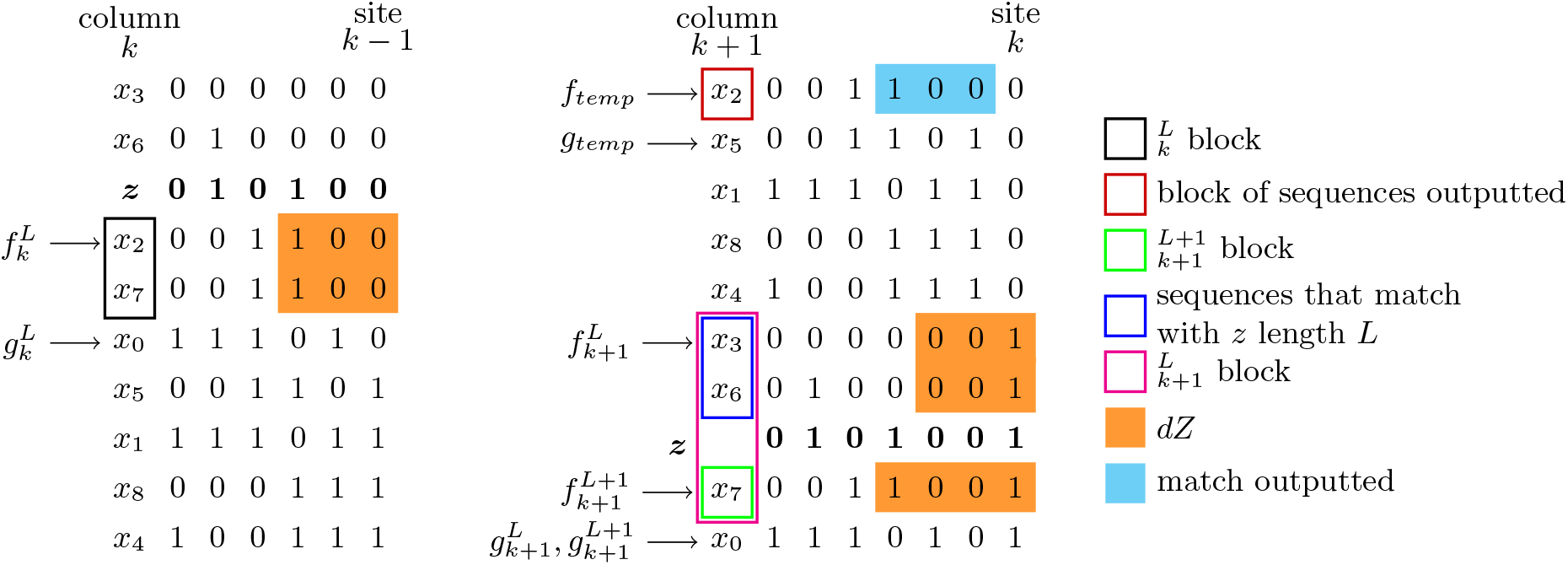
Computation of 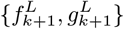 using 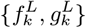 and extension function. *L* = 3. 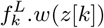 and 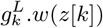 gives 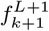 and 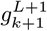. From there, the boundaries are expanded to include sequences that match with *z* length *L* until *k* + 1.The new boundaries are 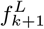 and 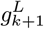. At the same time, 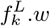(opposite of *z*[*k*]) and 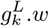(opposite of *z*[*k*]) is used to get *f*_*temp*_ and *g*_*temp*_. These mark the block of sequences that match with *z* length *L* or longer and the match ends at site *k*.

**Fig. 5.**
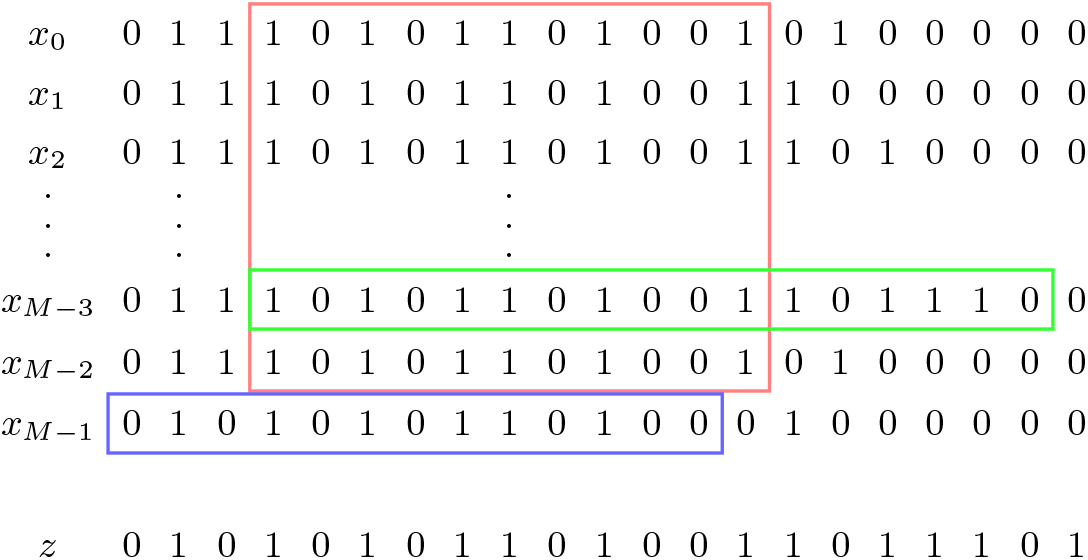
An example haplotype panel and *z* that causes Durbin’s algorithm 5 to run in *ω*(*N* + *c*) time.

The time complexity of this algorithm is easy to analyze. The Long Match Query algorithm (Algorithm 3) runs in worst case *O*(*N* + *c*) time. The virtual insertion portion of the algorithm runs in worst case *O*(*N*) because the haplotype panel and the *u* and *v* pointers are not updated. The query sweep loop has *N* iterations. All operations in one iteration of the query sweep loop take constant time except for the output while loop and the boundary expansion while loop. The output while loop will only output each match once, therefore the sum of all times it is entered in the algorithm is *c*, the number of matches found. The boundary expansion loop is entered once for every match that has length *L* (exactly) at some *k*. Every match will have length *L* exactly one time throughout the whole iteration of the algorithm, therefore the sum of all times the boundary expansion loop is entered is *c*. Therefore the algorithm runs in worst case *O*(*N* + *c*) time.

**Algorithm 3:**
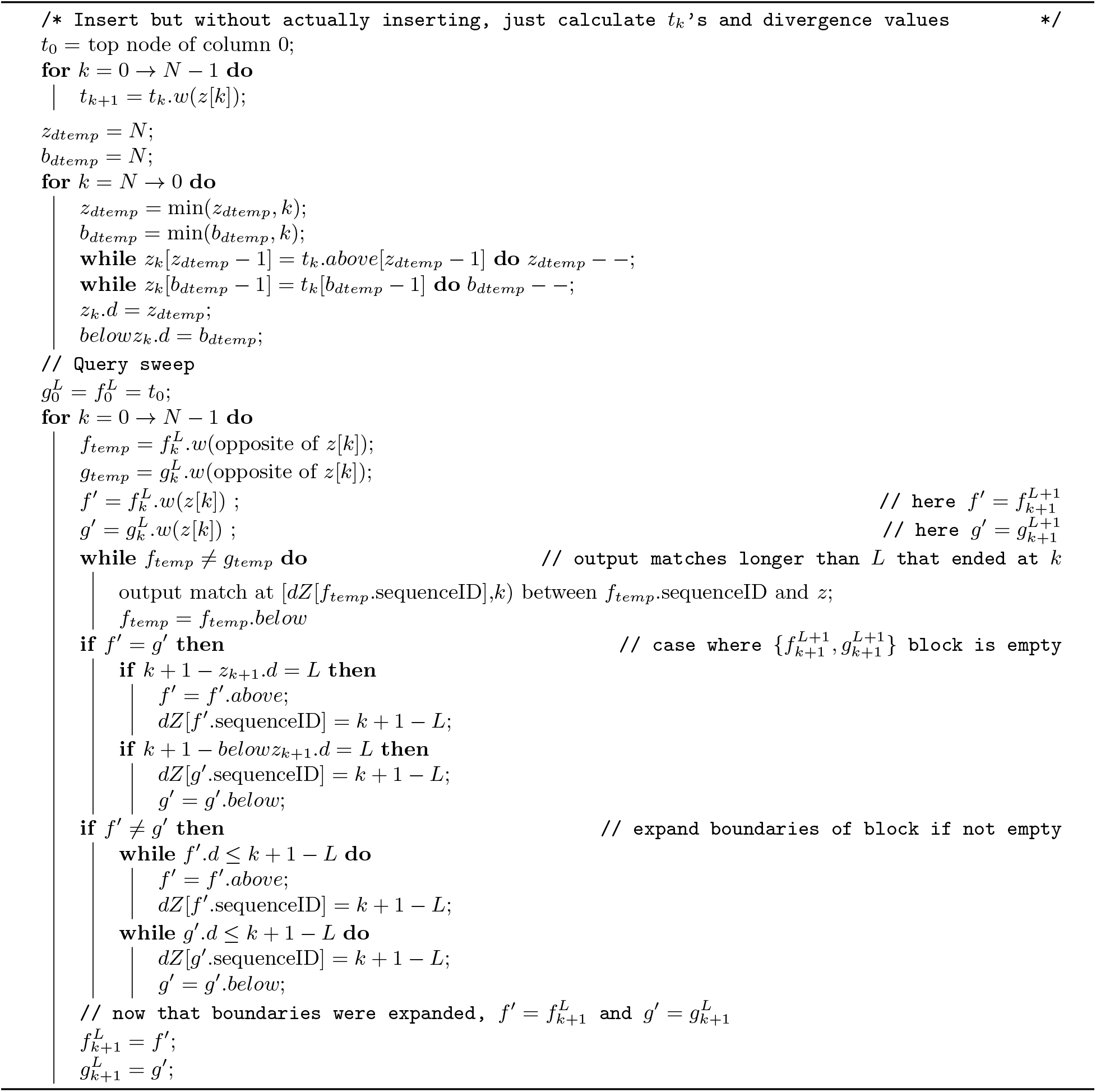
Long Match Query: Find long matches between query sequence *z* and sequences in the d-PBWT

### 2.6 Deletion

Deletion of a sequence from the d-PBWT is easy. If sequence *i* is to be deleted, sequence *x*_*M*−1_ needs to have the sequenceID of all its nodes changed from *M* − 1 to *i* so that the sequenceID definition is maintained after deletion of *x*_*i*_. Furthermore an array of pointers to the node in column 0 of each node needs to be kept so that the node of *x*_*i*_ in column 0 can be accessed in constant time. (Maintenance of this array is just an amortized constant time operation in the insertion and deletion algorithms.) The contiguous block of sequences above the node of *x*_*i*_ in column *k* neeeds to have their {*u* if *x*_*i*_[*k*] = 1, *v* otherwise} pointers updated. They are set to the value of the node below *x*_*i*_’s node. Lastly, the node of *x*_*i*_ in each column is deleted and the divergence of the node below it is updated. The whole algorithm can be done in one sweep. The time complexity of this algorithm is average case *O*(*N*). This is not worst case because of the update of the *u* and *v* pointers and haplotype panel. However, as stated before, update of the dynamic haplotype panel is amortized *O*(*N*) and the number of *u* and *v* pointers that will be updated per column will be a small constant on average. See Algorithm 4 in Appendix C.

### 2.7 Equivalence and Conversion between d-PBWT and PBWT

Equivalencies between data structures of PBWT and d-PBWT (Table 2) suggest that all construction algorithms and search algorithms can be translated between PBWT and d-PBWT with minimal changes. Moreover, d-PBWT data structure can be initialized by direct bulk conversion from an existing PBWT. Conversion of the d-PBWT to a PBWT in *O*(*MN*) time is trivial given its description. So is conversion of a PBWT to a d-PBWT. See Algorithms 5 and 6 in Appendix C.

Durbin’s Algorithms 1–5 can be implemented on the d-PBWT with a little modification. Furthermore, the pseudocode of our query algorithms are presented in the notation of d-PBWT, however, they can easily be applied to PBWT.

### 2.8 Single Sweep Long Match Query

While a Long Match Query algorithm that runs in worst case *O*(*N*) is an interesting theoretical development, an average case *O*(*N*) algorithm that only sweeps through the data once may be more useful for real world applications, particularly implementations that use memory mapping. The pseudocode for an average case *O*(*N*) Single Sweep Long Match Query algorithm (Algorithm 7) is provided in Appendix D. This is done using Durbin’s *e*_*k*_. Of course, the insertion algorithm can also be modified to run in a single sweep using *e*_*k*_.

## 3 Discussions

In this work, we developed the first dynamic PBWT data structure that allows efficient updating. When inserting or deleting a haplotype in a static PBWT panel, one has to reconstruct the entire PBWT panel in *O*(*MN*) time, while using dynamic PBWT, these can be achieved in Avg. *O*(*N*) time. In addition, we simplified and improved the PBWT query search algorithms (Durbin’s Algorithms 5 and L-PBWT-Query) in a worst case *O*(*N*) time and with no additional data structures. In doing so, we believe that we have brought the PBWT data structure closer to its full potential.

This work would enable efficient genealogical search in large databases. For example, large consumer-facing population databases hosting millions of individuals’ haplotypes typically have a constant burden of maintaining the population haplotype data structure in order to serve to report real-time genealogical search results. We believe that d-PBWT provides a practical solution for maintaining the population haplotype data structure. Our insertion and deletion algorithms can be implemented to handle high-volume updates in a real-time fashion. Meanwhile, the performance of genealogical search queries can be guaranteed by efficient long match query algorithms.

Notably, all three long match algorithms, including L-PBWT-query in Naseri *et al.* [5] and the Algorithms 3 and 7 presented here, achieve average case time complexity independent to database size. The only differences are their worst case time complexity, the number of sweeps required, and the memory needed for holding the additional auxiliary data structures. While in practice the optimal algorithm of choice may be a trade off of these and other factors, we believe Algorithm 7 provides a reasonable balance, as it only takes one sweep, with average linear time independent to panel size, and no additional memory for LEAP arrays.

Moreover, d-PBWT and our algorithms open new research avenues for developing efficient genotype imputation and phasing algorithms. Current practices of imputation and phasing are mainly based on a fixed reference panel. With d-PBWT, individual’s haplotypes in the reference panel can be iteratively refined, offering improved results.

## Acknowledgements

This work was supported by the US National Institutes of Health under grant number R01HG010086.

## Appendices

## A Time Complexity of Durbin’s Algorithm 5

Durbin claims “*The while loop in f′ or g′ is inevitable because it only takes as many iterations as there are matches to report the next time f′ = g′*” [2]. However, this is false. Figure 5 shows a possible haplotype panel that causes Durbin’s algorithm to run in *ω*(*N* + *c*) time. (All sequences between *x*_2_ and *x*_*M*−3_ match with *x*_0_ at [0, 15).) Therefore, at *k* = 13, Algorithm 5 will output *x*_*M*−1_ as a set maximal match at [0, 13) and the *f′* and *g′* loops will be entered to find the new block. The new block will have *M* − 1 sequences in it {*x*_0_ → *x_M−_*_2_}. However, only one of these sequences will be outputted as a set maximal match (*x*_*M*−3_ at [3, 20)). Therefore, the number of times the *f′* and *g′* while loops are entered is not bound by *c* (number of matches) and Algorithm 5 is not *O*(*N* + *c*). Of course, we have empirical evidence of the average case *O*(*N* + *c*) performance of Durbin’s Algorithm 5 [5].

## B Proofs

## B.1 Insertion

### Lemma 1.

*z*_*k*_.*d* ≤ *z*_*k*+1_*.d* ***and*** *z*_*k*_.*below.d* ≤ *z*_*k*+1_*.below.d*

*Proof.* If *z*_*k*+1_*.d > k*, then *z*_*k*_.*d* ≤ *z*_*k*+1_*.d* because *z*_*k*_.*d* ≤ *k*. Same for *z*_*k*+1_*.below.d* and *z*_*k*_.*below.d*.

The relative order of sequences that have the same value at site *k* is the same in column *k* and *k* + 1.

If *z*_*k*+1_*.d* ≤ *k* and *z*_*k*+1_*.below.d* ≤ *k*, then *z*_*k*_[*k*] = *z*_*k*+1_*.above*[*k*] = *z*_*k*+1_*.below*[*k*]. Therefore the relative order of *z*_*k*+1_, *z*_*k*+1_*.above,* and *z*_*k*+1_*.below* is the same in column *k* as it was in *k* + 1, i.e., *z*_*k*_.*above* is somewhere above *z*_*k*_ and *z*_*k*_.*below* is somewhere below *z*_*k*_.

If there is a sequence above *z*_*k*_ that matches longer than *z*_*k*+1_*.above*, it will be directly above *z* and *z*_*k*_.*d* < *z*_*k*+1_*.d*. If there is no sequence above *z*_*k*_ that matches longer than *z*_*k*+1_*.above*, then *z*_*k*+1_*.above* will be directly above *z*_*k*_ and *z*_*k*_.*d* = *z*_*k*+1_*.d*. Therefore *z*_*k*_.*d* ≤ *z*_*k*+1_*.d*

If there is a sequence below *z*_*k*_ that matches longer than *z*_*k*+1_*.below*, it will be directly below *z* and *z*_*k*_.*d* < *z*_*k*+1_*.d*. If there is no sequence below *z*_*k*_ that matches longer than *z*_*k*+1_*.below*, then *z*_*k*+1_*.below* will be directly below *z*_*k*_ and *z*_*k*_.*below.d* = *z*_*k*+1_*.below.d*. Therefore *z*_*k*_.*below.d* ≤ *z*_*k*+1_*.below.d*

## B.2 Set Maximal Match

### Lemma 2.

min(*z*_*k*_.*d, belowz*_*k*_.*d*) < min(*z*_*k*+1_*.d, belowz*_*k*+1_*.d*) *iff z’s locally maximal matches ending at k are set maximal*.

*Proof.* Assume min(*z*_*k*_.*d, belowz*_*k*_.*d*) < min(*z*_*k*+1_*.d, belowz*_*k*+1_*.d*) and

∃ sequence *s* and *d*_1_ < min(*z*_*k*_.*d, belowz*_*k*_.*d*) s.t. *s*[*d*_1_, *k*) = *z*[*d*_1_, *k*).

Then the local maximally matching sequence to *z* is not adjacent to it at *k* or the divergence values are incorrect (contradiction). Therefore there does not exist a sequence that has a match with *z* that extends this match to the left.

Assume min(*z*_*k*_.*d, belowz*_*k*_.*d*) < min(*z*_*k*+1_*.d, belowz*_*k*+1_*.d*) and

∃ sequence *s* and *k*_1_ > *k* s.t. *s*[min(*z*_*k*_.*d, belowz*_*k*_.*d*), *k*_1_) = *z*[min(*z*_*k*_.*d, belowz*_*k*_.*d*), *k*_1_).

Then min(*z*_*k*_.*d, belowz*_*k*_.*d*) = min(*z*_*k*+1_*.d, belowz*_*k*+1_*.d*) (contradiction). Therefore there does not exist a sequence that has a match with *z* that extends this match to the right.

Therefore there is no sequence that can extend this match and this match is locally maximal. So min(*z*_*k*_.*d, belowz*_*k*_.*d*) < min(*z*_*k*+1_*.d, belowz*_*k*+1_*.d*) ⇒ *z*’s locally maximal matches ending at *k* are set maximal.

Assume *z* has a set maximal match at [*d*_2_, *k*) and *d*_2_ = min(*z*_*k*_.*d, belowz*_*k*_.*d*) = min(*z*_*k*+1_*.d, belowz*_*k*+1_*.d*) then there is a match at [*d*_2_, *k* + 1) and the [*d*_2_, *k*) match can be extended. Therefore it is not set maximal (contradiction). We have already shown that the divergence at *k* ≤ divergence at *k* + 1 in **Lemma 1**. Therefore *z*’s locally maximal matches ending at *k* are set maximal ⇒ min(*z*_*k*_.*d, belowz*_*k*_.*d*) < min(*z*_*k*+1_*.d, belowz*_*k*+1_*.d*).

Therefore min(*z*_*k*_.*d, belowz*_*k*_.*d*) < min(*z*_*k*+1_*.d, belowz*_*k*+1_*.d*) ⟺ *z*’s locally maximal matches ending at *k* are set maximal.

## B.3 Long Match

### Lemma 3.

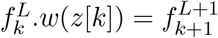 *and* 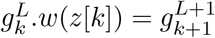.

*Proof.* All the sequences that match with *z* length *L* + 1 or longer until *k* + 1 all match with *z* length *L* or longer until *k*,i.e., the set of sequences that match with *z* length *L* + 1 until *k* + 1 is a subset of the set of sequences that match with *z* length *L* or longer until *k*. Specifically, it is the subset that has the same value at *k* as *z*.

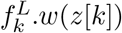 gives us the (node in column *k* + 1 of the) first sequence after 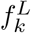 (inclusive) that has *z*[*k*] at position *k*. This is the first sequence in the 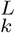 block that has *z*[*k*] at *k*. Since relative order of sequences with the same value is preserved, this will be the first sequence of the 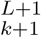 block.

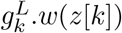 gives us the (node in column *k* + 1 of the) first sequence after 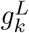 (inclusive) that has *z*[*k*] at position *k*. This is the first sequence outside of the 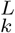 block that has *z*[*k*] at *k*. Since relative order of sequences with the same value is preserved, this will be the first sequence after the 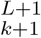 block.

Therefore 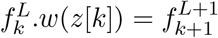 **and** 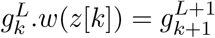

## C Deletion and Conversion Pseudocode

**Algorithm 4:**
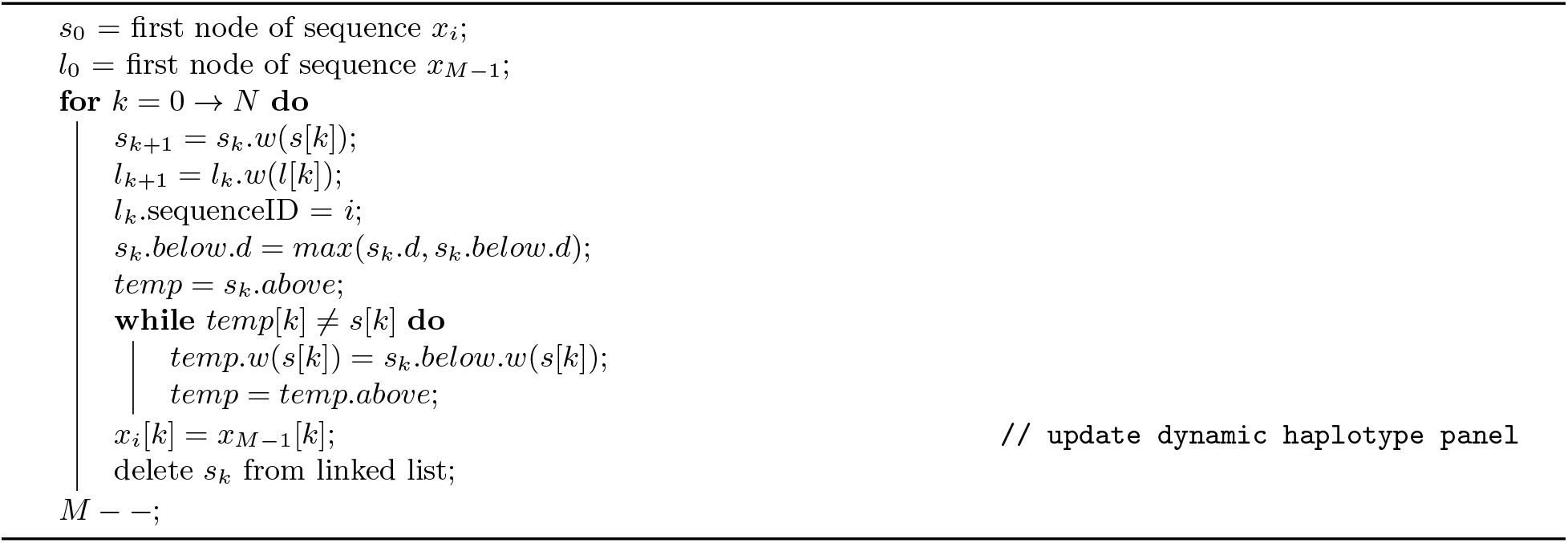
Deletion: Delete sequence *x*_*i*_ from d-PBWT

**Algorithm 5:**
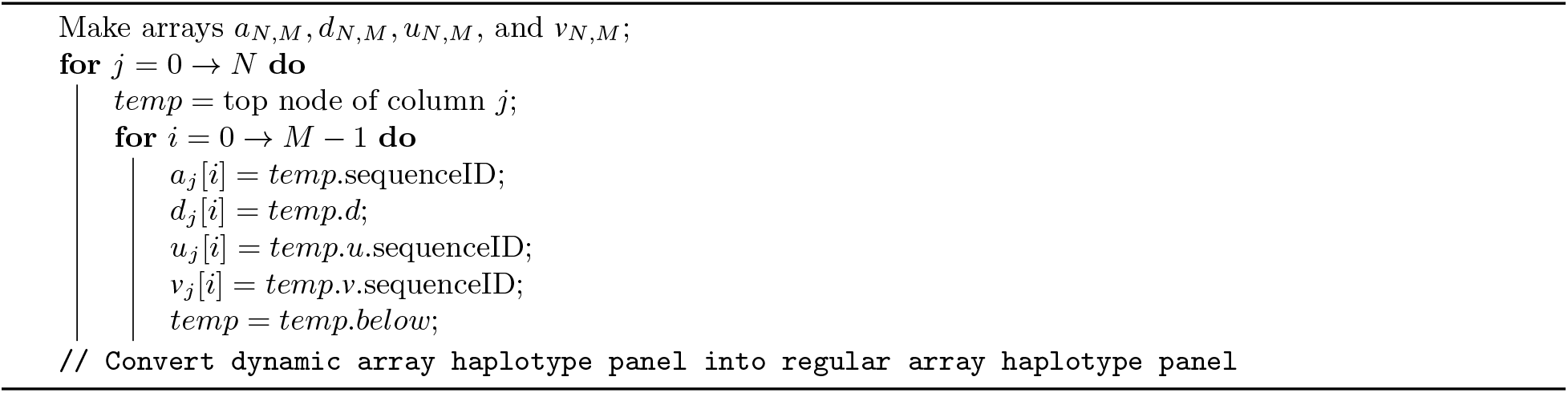
Conversion1: Converts d-PBWT into an array PBWT

**Algorithm 6:**
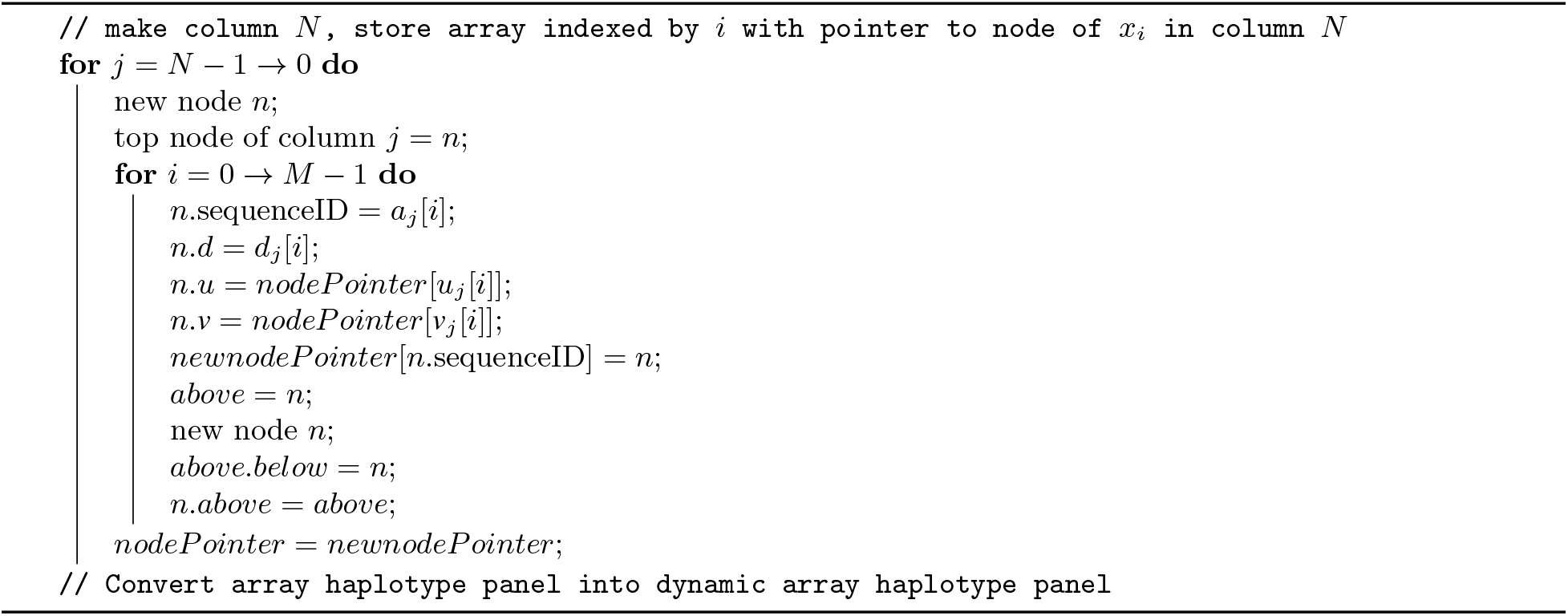
Conversion2: Converts PBWT into d-PBWT

## D Single Sweep Long Match Query Pseudocode

**Algorithm 7:**
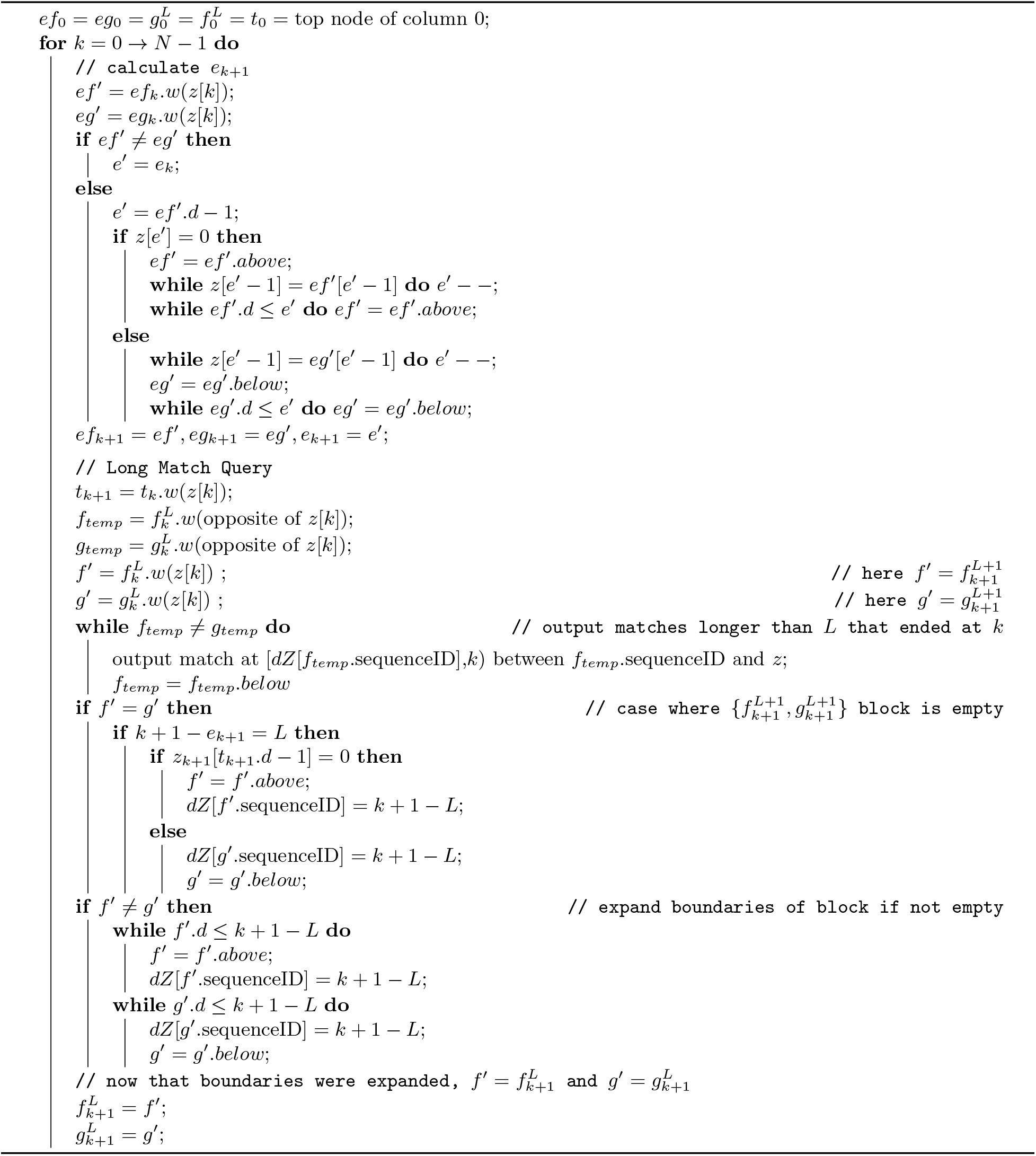
Single Sweep Long Match Query: Find long matches between *z* and sequences in d-PBWT in average case *O*(*N*) time and one sweep

## Footnotes

3 While *column* is an array-biased term, we abuse it for convenience of corresponding back to array-based PBWT.

4 It is OK to use array for indexing columns as long as the sites of a genome are stable. However, it may be possible to extend the columns to be non-linearly sorted, as in *variant graph* [3].

5 Durbin says in section 2.5: “*This is because population genetic structure means that there is local correlation in values due to linkage disequilibrium, which means that haplotypes with similar prefixes in the sort order will tend to have the same allele values at the next position, giving rise to long runs of identical values in the y array*” [2].

